# Endometrial decidualization status modulates endometrial perivascular complexity and trophoblast outgrowth in gelatin hydrogels

**DOI:** 10.1101/2022.11.08.515680

**Authors:** Samantha G. Zambuto, Hannah Theriault, Ishita Jain, Cody O. Crosby, Ioana Pintescu, Noah Chiou, Janet Zoldan, Gregory H. Underhill, Kathryn B.H. Clancy, Brendan A.C. Harley

## Abstract

The endometrium undergoes rapid cycles of vascular growth, remodeling, and breakdown during the menstrual cycle and pregnancy. Decidualization is an endometrial differentiation process driven by steroidal sex hormones that is critical for blastocyst-uterine interfacing and blastocyst implantation. Certain pregnancy disorders may be linked to decidualization processes. However, much remains unknown regarding the role of decidualization and reciprocal trophoblast-endometrial interactions on endometrial angiogenesis and trophoblast invasion. Here, we report an artificial endometrial perivascular niche embedded in gelatin methacrylol hydrogels that displays morphological and functional patterns of decidualization. We show vessel complexity and soluble factor secretion are sensitive to decidualization and affect trophoblast motility. Importantly, we demonstrate the engineered perivascular niche can be combined with epithelial cultures to form a stratified endometrial model. This artificial perivascular niche provides a well-characterized platform to investigate dynamic changes in angiogenesis in response to pathological and physiological endometrial states.

**Teaser:** We describe an endometrial vessel model to understand endometrial vasculature in the menstrual cycle and pregnancy.

## Introduction

To support tissue regrowth during the menstrual cycle and significant remodeling during pregnancy, the lining of the uterus referred to as the endometrium undergoes rapid cycles of vascular growth, remodeling, and breakdown. The endometrium is one of the only adult human tissues to undergo non-pathological angiogenesis (*1*): endothelial cell-driven development of new vessels from existing blood vessels via elongation, intussusception, or sprouting (*1, 2*). During the menstrual cycle, angiogenesis occurs during three distinct phases: the proliferative phase, the secretory phase, and the menstrual phase (*1, 3*). The proliferative phase is characterized by initiation of vessel growth (*1, 3*). In the secretory phase, the vessels continue to grow by branching, lengthening, and maturing (*1, 3*). Menstruation then induces vessel degeneration and endometrial shedding (*3*). During menstruation, angiogenic processes also work to repair the superficial layer of the basal endometrium in preparation for the subsequent menstrual cycle (*1*). During pregnancy, rapid angiogenesis also occurs immediately after trophoblast-uterine interfacing and prior to the formation of a placenta during pregnancy due to significant demands to support a growing placenta and eventual fetus (*3*). Coordinated angiogenesis provides essential early support necessary to maintain these structures and their development (*3*).

Sex steroid hormones progesterone and estrogen modulate endometrial angiogenesis and remodeling (*3*). Progesterone controls vessel elongation, growth and coiling of spiral arterioles, and maturation of the subepithelial capillary plexus whereas estrogen plays a key role in concert with the VEGF (vascular endothelial growth factor) family to control vascular remodeling (*3*). These hormones also orchestrate a differentiation process known as decidualization. Decidualization is the process by which the endometrium prepares for a potential pregnancy by thickening and enhancing the tissue matrix for the incoming blastocyst (*3*). During decidualization, vessels sprout and lengthen, the surface area of spiral arterioles increases, uterine glands undergo secretory transformation, and specialized uterine natural killer cells increase in number (*3*). Successful endometrial decidualization enables the endometrium to enter a period of receptivity that occurs during the late secretory phase of the menstrual cycle (*3*). During this window, the blastocyst can attach to the endometrial epithelium and subsequently invade into the underlying stroma and vasculature (*3*). Crosstalk between endometrial cells and trophoblast cells from the invading blastocyst are believed to modulate processes of invasion and dynamic vascular remodeling (*3*). Endometrial decidualization is critical here, with reduced decidualization linked to a variety of pregnancy disorders such as preeclampsia, fetal growth restriction, and infertility (*3*). For example, in the hypertensive pregnancy disorder preeclampsia, studies have demonstrated that in patients with severe preeclampsia, reduced decidualization subsequently led to impaired trophoblast invasion (*4*). However, ethical considerations and technical limitations limit the ability to study these processes *in vivo*. As a result, much remains unknown regarding the role of sex steroid hormones on endometrial angiogenesis and vice versa.

Few models of endometrial vasculature exist. Most models of the endometrial vasculature either cannot recapitulate tissue biophysical properties (stiffness), do not use relevant human cell types in heterogenous cell cultures, or cannot be cultured long term (20+ days) days (*5–9*). Engineered vasculature models have recently demonstrated promising results for mimicking vasculature *in vitro* for a wide range of applications. For example, Offendu et al. and Haase et al. used perfusable vasculature in microphysiological microfluidic devices for quantifying transport through the endothelium and flow-mediated vessel remodeling (*10–12*). Such model systems have also been used to study pregnancy-related vascular disorders, including placental vasculopathies (*11*). Engineered vasculature models can support three-dimensional culture of heterogeneous cell populations in biomaterials that can recapitulate tissue biophysical properties. Gelatin is an attractive platform for these types of studies for a variety of reasons. As a natural polymer derived from collagen, it contains cell adhesion and degradation sites which allow for matrix remodeling by cells. Functionalization of gelatin by adding methacrylate groups to its amine-containing side groups results in the synthesis of gelatin methacrylol, GelMA, which offers enhanced mechanical features that can be tuned to mimic *in vivo* biophysical properties (*13*). Previous studies from our group and others have demonstrated that vessel networks can be cultured in GelMA hydrogels by encapsulating co-cultures of endothelial and stromal cells (*13, 14*).

Here, we develop and characterize models of an artificial endometrial perivascular niche embedded in GelMA hydrogels that display morphological and functional patterns of decidualization. We quantify shifts in vessel network complexity, analyze soluble factor secretion of perivascular cultures, and assess matrix remodeling via basement membrane protein deposition and tight junction formation. Subsequently, we examine bidirectional communication, notably the role of trophoblast secreted factors on perivascular network remodeling and the role of perivascular secreted factors on trophoblast motility. Finally, we demonstrate a three-dimensional, stratified model of the endometrium consisting of an epithelial culture overlaying an embedded perivascular niche. This artificial perivascular niche replicates aspect of the *in vivo* perivascular environment and can serve as a platform to study endometrial angiogenesis in a variety of endometrial states.

## Results

### Human endometrial microvascular endothelial cells (HEMEC) demonstrate angiogenic potential

We first assessed phenotypic markers and the ability to form vessel structures of human endometrial microvascular endothelial cells (HEMECs). HEMEC plated on 2D plates expressed CD31 (Supplemental Fig. 1A) and von Willebrand factor (Supplemental Fig. 1B). HEMECs plated on Matrigel transiently formed tubes that fell apart in less than 24 hours (Supplemental Fig. 1B). Taken together, these results suggest HEMECs express characteristic endothelial cell markers and they have the potential to form vessel-like structures *in vitro*.

### Endometrial perivascular niche hydrogels can be cultured long term

The longevity and stability of engineered endometrial perivascular niches (PVNs) were tracked for up to 28 days of culture *in vitro*. All PVNs contained a constant number of endothelial cells (500,000 cells/mL) but variable number of stromal cells at ratios of 2:1 (250,000 stromal cells/mL), 1:1 (500,000 stromal cells/mL), and 1:2 (1,000,000 stromal cells/mL) endothelial:stromal cells. Notably, the ability to form stable perivascular models was tightly tied to the ratio of perivascular cells. For the 1:1 endothelial:stromal laden hydrogels, total network length, branches, and vessels all increased from days 7 to 14 while branch length decreased (Fig. 2B-E); however, by day 21 all 1:1 hydrogels had disintegrated. Increasing the number of stromal cells (1:2 ratio) did not improve perivascular niche stability. Here, total network length, branches, and vessels all increased from days 7 to 14 but then decreased from days 14 to 21 (Fig. 2B-E) while branch length decreased between days 7 and 14 but increased between days 14 and 21. Although the 2:1 endothelial:stromal ratio PNV initially contained the least number of stromal cells, it was the most stable over time. The 2:1 endothelial:stromal ratio showed increasing metrics of total network length, branches, and vessels from days 7 to 21 while branch length appeared to remain consistent across all days with only a slight decrease over time (Fig. 2B-E). No culture remained stable through 28 days, with hydrogel degradation being the primary limitation of culture stability over long time periods.

**Fig. 1.**
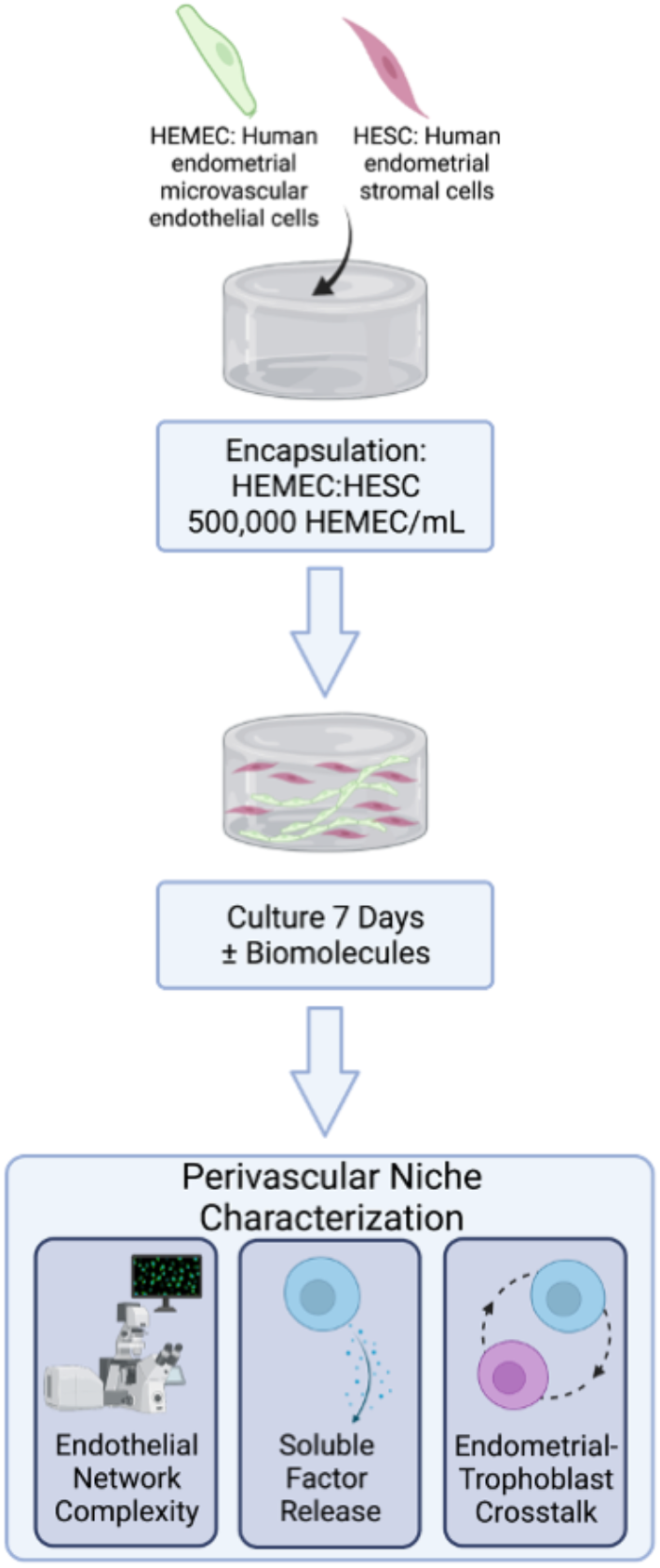
Development and characterization of an artificial endometrial perivascular niche model. Encapsulated endometrial endothelial and stromal cells are co-cultured in methacrylamide-functionalized gelatin hydrogels for 7 days and are subsequently analyzed for vessel network complexity, soluble factor secretion, and matrix remodeling. Created with Biorender.com.

**Fig. 2.**
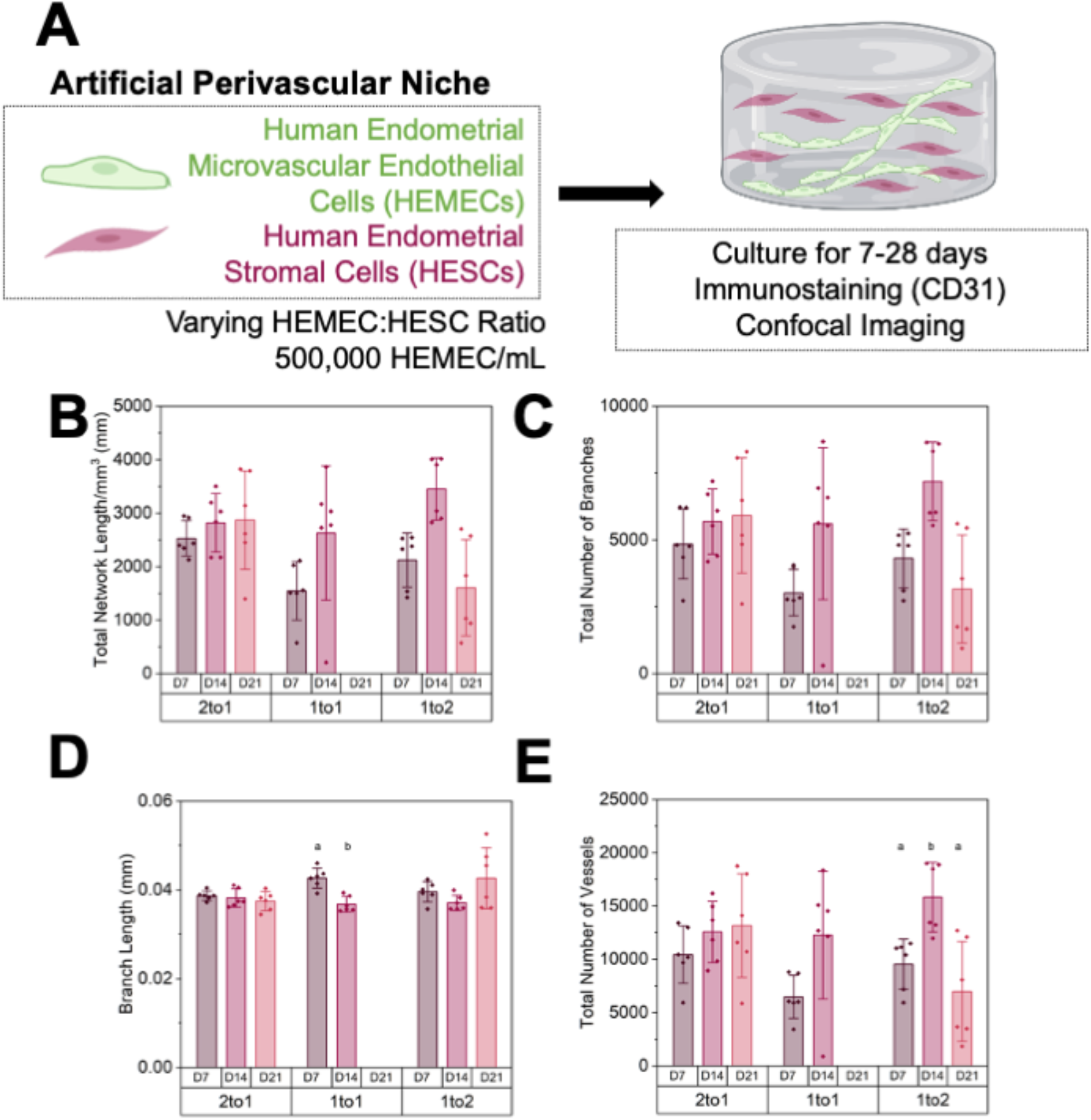
28-day culture of an artificial endometrial perivascular niche. **(A)** Experimental summary. **(B)** Quantification of total vessel length per mm^3^, **(C)** total number of branches, **(D)** average branch length, and **(E)** total number of vessels at days 7, 14, and 21 (n=6 hydrogels per condition; 3 ROI imaged per gel and averaged) of varying endothelial to stromal cell ratios. Groups with different letters are statistically significantly different from each other. Data presented as mean ± standard deviation. Created with Biorender.com.

### Ratio of endothelial to stromal cells affects vessel complexity more so than addition of VEGF

Soluble VEGF was added to culture media to determine if additional exogenous proangiogenic factors would affect vessel complexity. We observed no significant differences in trends of metrics of perivascular network complexity as a result of VEGF inclusion (Supplemental Fig. 2) and concluded that the endometrial PVNs do not require supplementation with additional proangiogenic factors. Based on these results and the results of the time course experiment, the 2:1 endothelial:stromal cell ratio PVN was used for all subsequent experiments because this condition resulted in the most consistent, stable networks over time.

### PVN cultures deposit laminin and express tight junction marker ZO-1

Metrics of basement membrane protein deposition and expression of tight junction proteins were subsequently assessed for engineered endometrial PVNs. Immunofluorescent staining demonstrated laminin, a common basement membrane protein, deposition by both cell types (Supplemental Fig. 3B). Additionally, ZO-1 expression was observed by day 7 of culture, suggesting formation of tight junction in our PVNs (Supplemental Fig. 3C). We deployed a spatial analysis tool to quantify the degree of overlap between endometrial vessels (CD31+) and laminin/ZO-1 (Supplemental Fig. 3D-E). We calculated the average pixel intensity for both stains and calculated the degree of overlap by multiplying the binarized matrix of vessels by the binarized matrix of the proteins. Laminin/CD31 and ZO-1/CD31 both displayed approximately 30% signal overlap, suggesting that not only are laminin and ZO-1 expressed in engineered endometrial PVNs but also they are expressed in close proximity to the endometrial perivascular networks formed within gelatin hydrogels.

### Endometrial stromal cell decidualization status modulates endometrial PVN complexity

Decidualization of the engineered endometrial perivascular networks was assessed in response to two decidualization protocols previously reported for two-dimensional cell culture (*15–20*): exogenous addition of 1 μM synthetic progestin medroxyprogesterone acetate (MPA), 0.5 mM 8-bromodenosine 3’,5’-cyclic monophosphate (8-Br-cAMP) or 0.5 mM dibutyryl cyclic AMP (dcAMP) + 10 nM estradiol (E2) + 100 nM progesterone (P4). Morphological changes to the perivascular networks were first compared to a control condition (no decidualization hormones). Total network length, branches, and vessels increased in response to MPA decidualization condition compared to control (Fig. 3B-E). Branch length was reduced for the MPA and P4 groups in comparison to control (Fig. 3D). Total number of vessels increased for both decidualization conditions, with the MPA decidualization condition having the largest total number of vessels (Fig. 3E). Total network length and branches were not different between control and P4 decidualization conditions (Fig. 3B-C). Total network length and average branch length were not different between MPA and P4 groups but total branches were increased in the MPA group compared to the P4 group (Fig. 3B-E). Taken together, these results suggest decidualization status as well as choice of decidualization hormone cocktail strongly influence vessel network complexity in gelatin hydrogels, with decidualization broadly resulting in a denser network of smaller endothelial cell networks (increased number of shorter branches) and the strongest effect seen for decidualization with synthetic progesterone.

**Figure 3.**
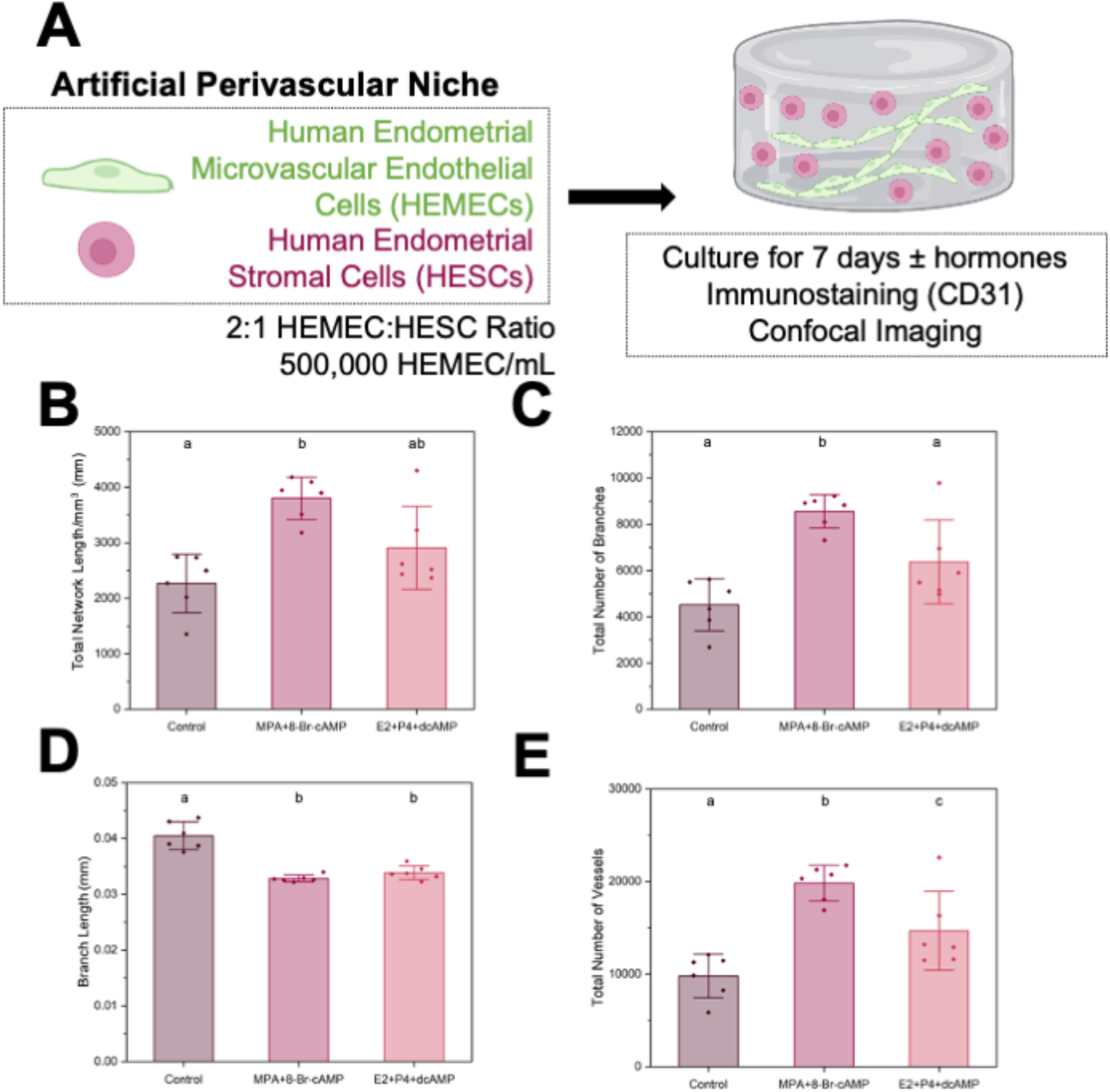
Stromal cell decidualization in an artificial endometrial perivascular niche. **(A)** Experimental summary. **(B)** Quantification of total vessel length per mm^3^, **(C)** total number of branches, **(D)** average branch length, and **(E)** total number of vessels for control and decidualized samples (n=6 hydrogels per condition; 3 ROI imaged per gel and averaged). Two decidualization conditions were tested. Control condition contained no added decidualization hormones. Groups with different letters are statistically significantly different from each other. Data presented as mean ± standard deviation. MPA: medroxyprogesterone acetate, Br-cAMP: bromoadenosine cyclic AMP, E2: estradiol, P4: progesterone, dcAMP: dibutyryl cyclic AMP. Created with Biorender.com.

### Decidualization status strongly influences endometrial PVN secretome

We subsequently examined biomolecular consequences of decidualization (control, decidualized-MPA, decidualized-P4) of perivascular networks via a cytokine array (quantifying mean pixel density of cytokine spots normalized to a positive control, Fig. 4B). We first compared secretion of characteristic markers of decidualization prolactin and IGFBP-1 (insulin-like growth factor binding protein 1) (Fig. 4C). Prolactin and IGFBP-1 secretion increased for decidualized conditions compared to control, strongly indicating that stromal cells were decidualized in the presence of decidualization hormones. Subsequently, we performed statistical analysis across 55 human angiogenesis associated proteins (statistical tests and p-values: Supplemental Figures 4-5). We identified 14 proteins whose expression was significantly altered by PVN decidualization: activin A, angiogenin, angiopoietin-1, amphiregulin, endoglin, endostatin/collagen XVIII, endothelin-1, FGF-1 (FGF acidic), IGFBP-2, Pentraxin 3 (PTX3), PDGF-AA, Platelet Factor 4 (PF4), Prolactin, and Serpin F1 (Fig. 4D; Table 1). We then used STRING to generate a network summary of predicted protein associations between the 14 proteins (Fig. 4E). The STRING network contained 14 nodes, 25 edges, 3.57 average node degree with an average local clustering coefficient of 0.495 and PPI enrichment p-value < 1.0×10^−16^, and identified interactions between these factors, except for PF4, Prolactin, and Amphiregulin. Gene ontology analysis suggests these cytokines play critical roles in blood vessel development and branching, regulating endothelial cell proliferation, and epithelium branch elongation, with many of these cytokines associated with the basement membrane, ECM, extracellular space, and cytoplasmic vesicles.

**Figure 4.**
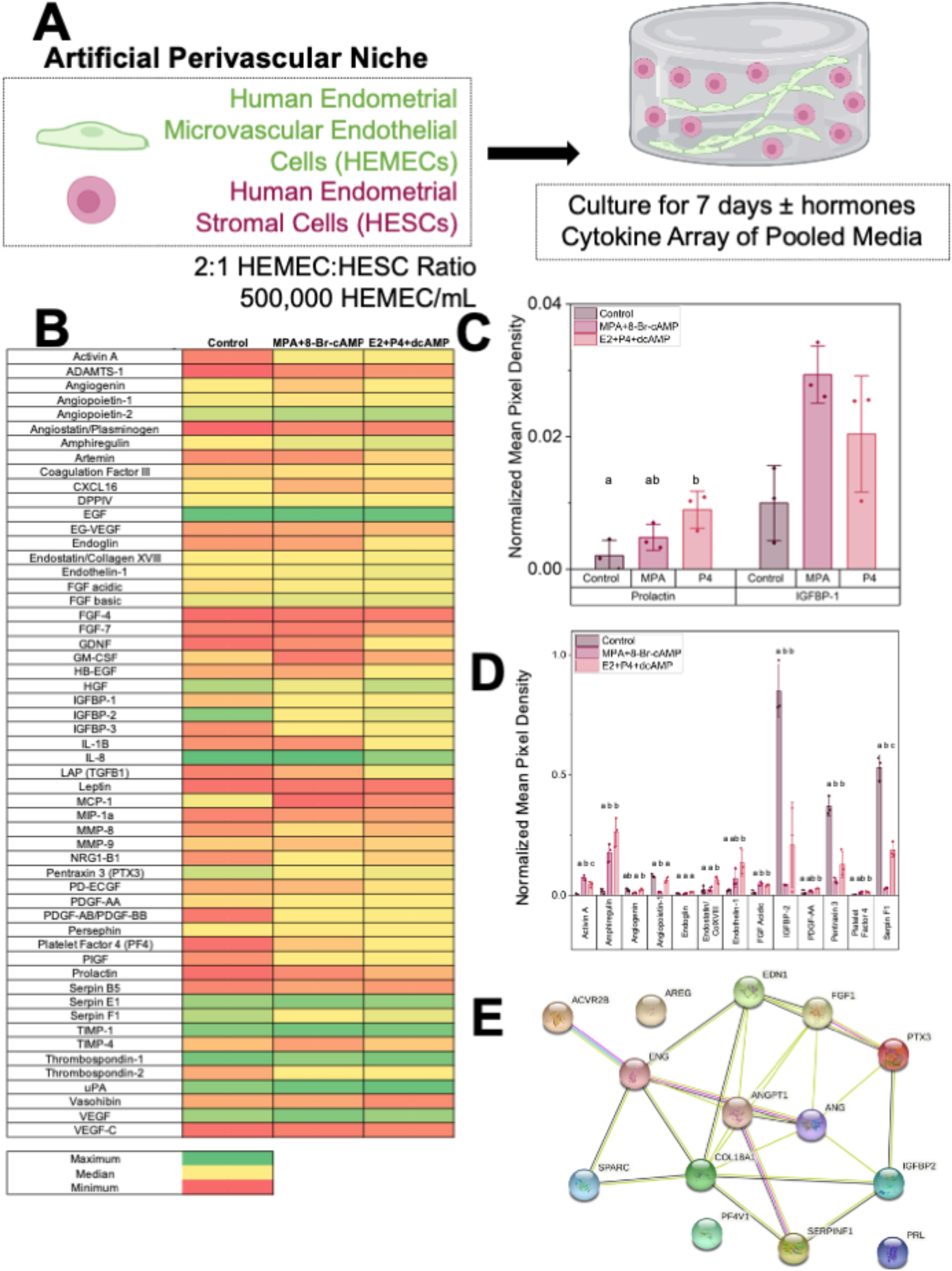
Cytokine secretion across control and decidualized artificial perivascular niche samples. **(A)** Experimental summary. **(B)** Cytokine array normalized mean pixel density results. **(C)** Normalized mean pixel density for characteristic decidual proteins Prolactin and IGFBP-1 across groups. **(D)** Normalized mean pixel density for statistically significantly different cytokines across groups. Groups with different letters are statistically significantly different from each other. Data presented as mean ± standard deviation. MPA: medroxyprogesterone acetate, Br-cAMP: bromoadenosine cyclic AMP, E2: estradiol, P4: progesterone, dcAMP: dibutyryl cyclic AMP. N=3 hydrogels per condition. **(E)** STRING analysis of statistically significantly different cytokines in homo sapiens. Red line-fusion evidence. Green line-neighborhood evidence. Blue line-concurrence evidence. Purple line-experimental evidence. Yellow line-textmining evidence. Light blue line-database evidence. Black line-coexpression evidence. COL18A1-Collagen alpha-1 (XVIII). ANGPT1-Angiopoetin-1. FGF1-Fibroblast growth factor 1. AREG-Amphiregulin. PTX3-Pentaxin-related protein 3. EDN1-Endothelin-1. ENG-Endoglin. ANG-Angiogenin. PRL-Prolactin. ACVR2B-Activin receptor type-2B. PF4V1-Platelet factor 4 variant. IGFBP-2-Insulin-like growth factor binding protein 2. FGF1-Fibroblast growth factor 1. SERPINF1-Pigment epithelium derived factor. Created with Biorender.com.

**Table 1.**
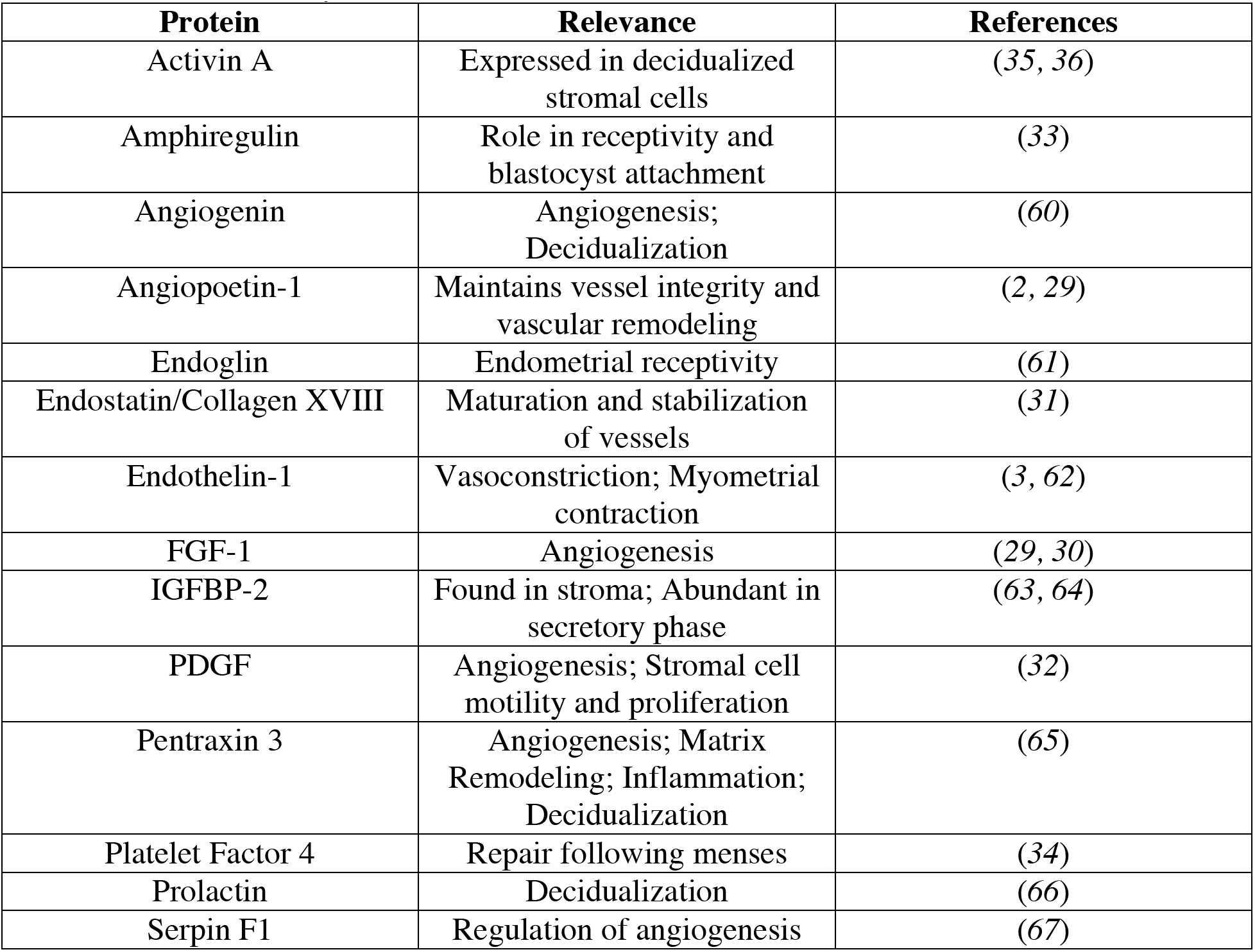
Relevance of cytokines to endometrial function.

### Perivascular niche secreted factors influence trophoblast invasion, but trophoblast secreted factors do not affect vessel network complexity

We then examined reciprocal crosstalk between endometrial perivascular cells and trophoblast cells. We first examined how perivascular niche secreted factors affect trophoblast motility. We quantified Swan71 trophoblast outgrowth area for 3 days in response to conditioned media collected from control or decidualized perivascular niche hydrogel cultures using a previously described cell spheroid assay that quantifies total outgrowth area of spheroid (Fig. 5). We collected conditioned media from non-decidualized (CM non-decidualized) or decidualized (CM MPA or CM P4) perivascular niches and compared trophoblast invasion against two unique media controls (control or media control). We defined the control condition as conventional trophoblast growth medium. The media control condition was a 1:1 mixture of trophoblast and PVN growth medium. This second control was used to replicate the media composition of the experimental groups, which contained a 1:1 mixture of trophoblast media and conditioned media collected from PVN cultures.

**Figure 5.**
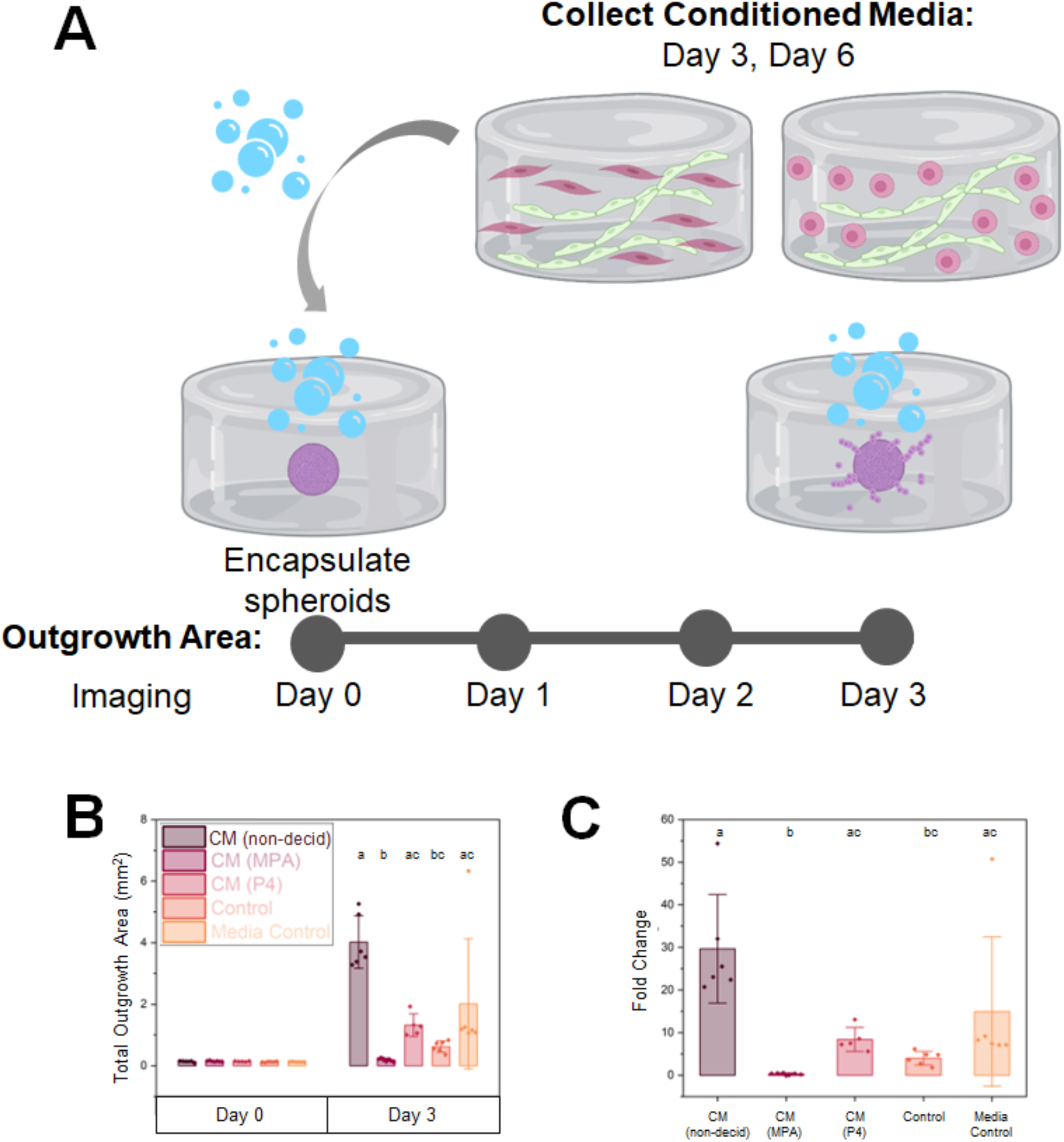
The effects of perivascular conditioned media on trophoblast motility. **(A)** Experimental summary. **(B)** Quantification of total outgrowth area (mm^2^) and **(C)** fold change in outgrowth area at Day 3 compared to Day 0 (encapsulation). Groups with different letters are statistically significantly different from each other. Data presented as mean ± standard deviation. CM: Conditioned Media, Non-decid: not decidualized (no hormones), MPA: medroxyprogesterone acetate, P4: progesterone. Created with Biorender.com.

By day 3, we observed significant (*p*=3.8×10^−5^) differences in outgrowth area and fold change in outgrowth area (outgrowth at day 3 normalized to the same spheroid at day 0; Fig. 5C). Dunn’s post hoc analysis revealed four significant differences between outgrowth area groups: conditioned media (control-not decidualized) and conditioned media (MPA-decidualized) (*p*=4.35×10^−5^), conditioned media (control-not decidualized) and control (*p*=0.016), conditioned media (MPA-decidualized) and conditioned media (P4-decidualized) (*p*=0.036), and conditioned media (MPA-decidualized) and media control (*p*=0.011). Fold change was then calculated by normalizing initial spheroid outgrowth area to outgrowth area on day 3. The same trend observed in outgrowth area was also observed in fold change in outgrowth area, with *p*=3.69×10^−5^, *p*=0.016, *p*=0.046, and *p*=0.011, respectively. These findings suggest that soluble factors from perivascular cultures increase trophoblast outgrowth for non-decidualized and decidualized conditions with P4 compared to the control condition. However, trophoblast outgrowth was decreased in decidualized conditions with MPA compared to the control condition. These findings demonstrate that not only perivascular decidualization status but also mode of decidualization affect trophoblast outgrowth.

We then assessed whether and how trophoblast secreted factors influenced perivascular niche complexity. We quantified metrics of vessel network complexity in the presence and absence of conditioned medium from Swan71 trophoblast cells (Fig. 6). The control condition contained PVN growth medium, and the media control condition was a mixture of trophoblast and PVN growth medium consistent with the conditioned medium conditions. We observed no differences in total network length/mm^3^, total number of branches, and total number of vessels in the presence of conditioned media from Swan71 trophoblast cells (Fig. 6B,C,E). Although we did observe branch length decreased between the media control (50:50 media ratio) and control (Swan71 invasion media) conditions (Fig. 6D), this was likely due to increased serum content not depleted by cells.

**Figure 6.**
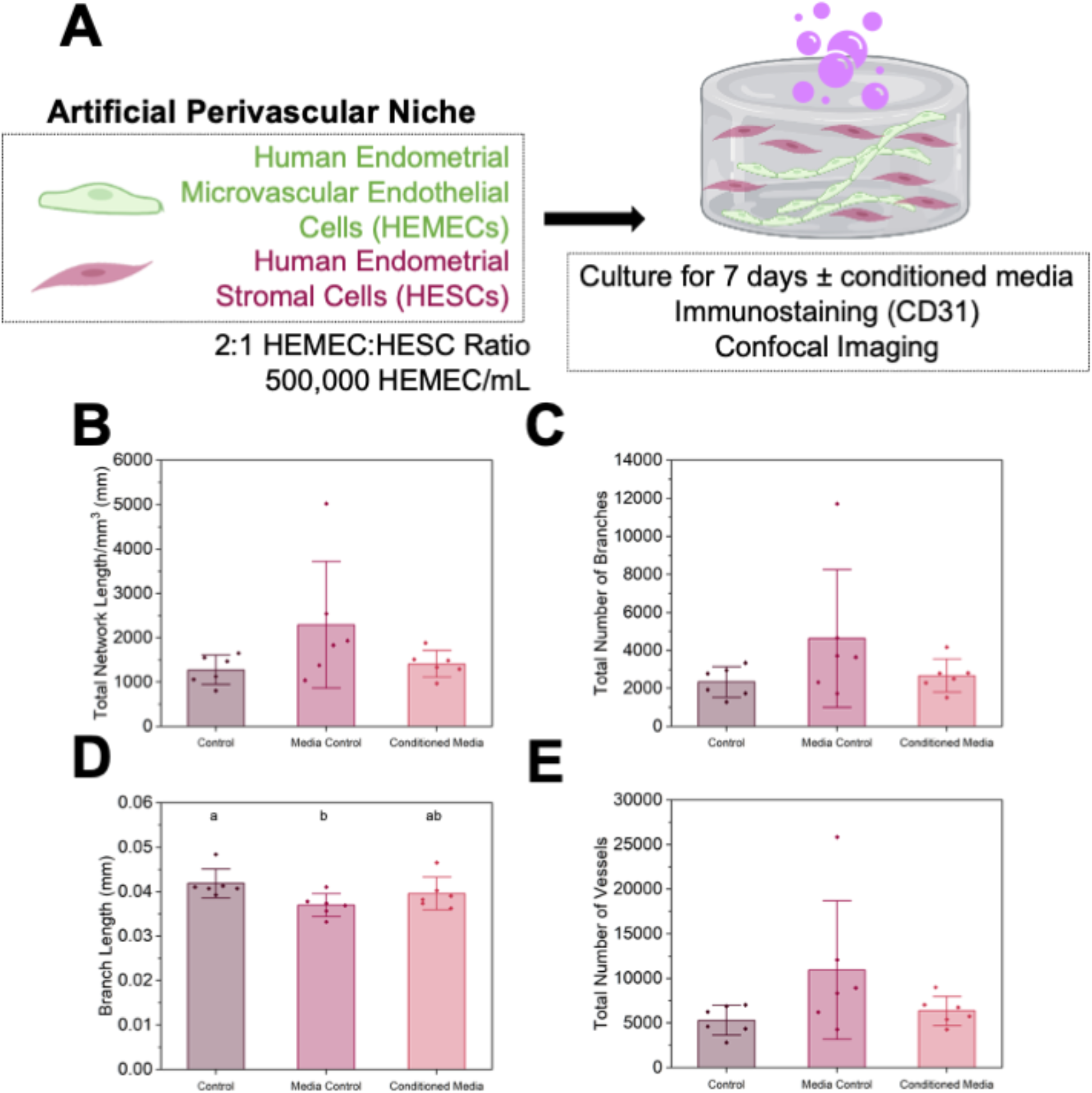
Effects of Swan71 trophoblast conditioned medium on endometrial perivascular niche complexity. **(A)** Experimental summary. **(B)** Quantification of total vessel length per mm^3^, **(C)** total number of branches, **(D)**average branch length, and **(E)** total number of vessels for control, media control, and conditioned media samples (n=6 hydrogels per condition; 3 ROI imaged per gel and averaged). Groups with different letters are statistically significantly different from each other. Data presented as mean ± standard deviation. Created with Biorender.com.

### Stratified endometrial epithelial culture overlaying a perivascular niche

Finally, we fabricated an endometrial triculture to replicate features of the stratified endometrium *in vivo* by seeding primary endometrial epithelial cells on top of the 3D hydrogel perivascular culture (Fig. 7). We observed two stratified components: an epithelial layer overlying an embedded perivascular culture (Fig. 7B). We subsequently examined epithelial cell morphology and phenotype via immunohistochemistry for CK18, a marker of epithelial cell attachment (Fig. 7C). We observed regions of epithelial monolayers that positively express CK18, suggesting the epithelial monolayer is attached to the underlying perivascular hydrogel.

**Figure 7.**
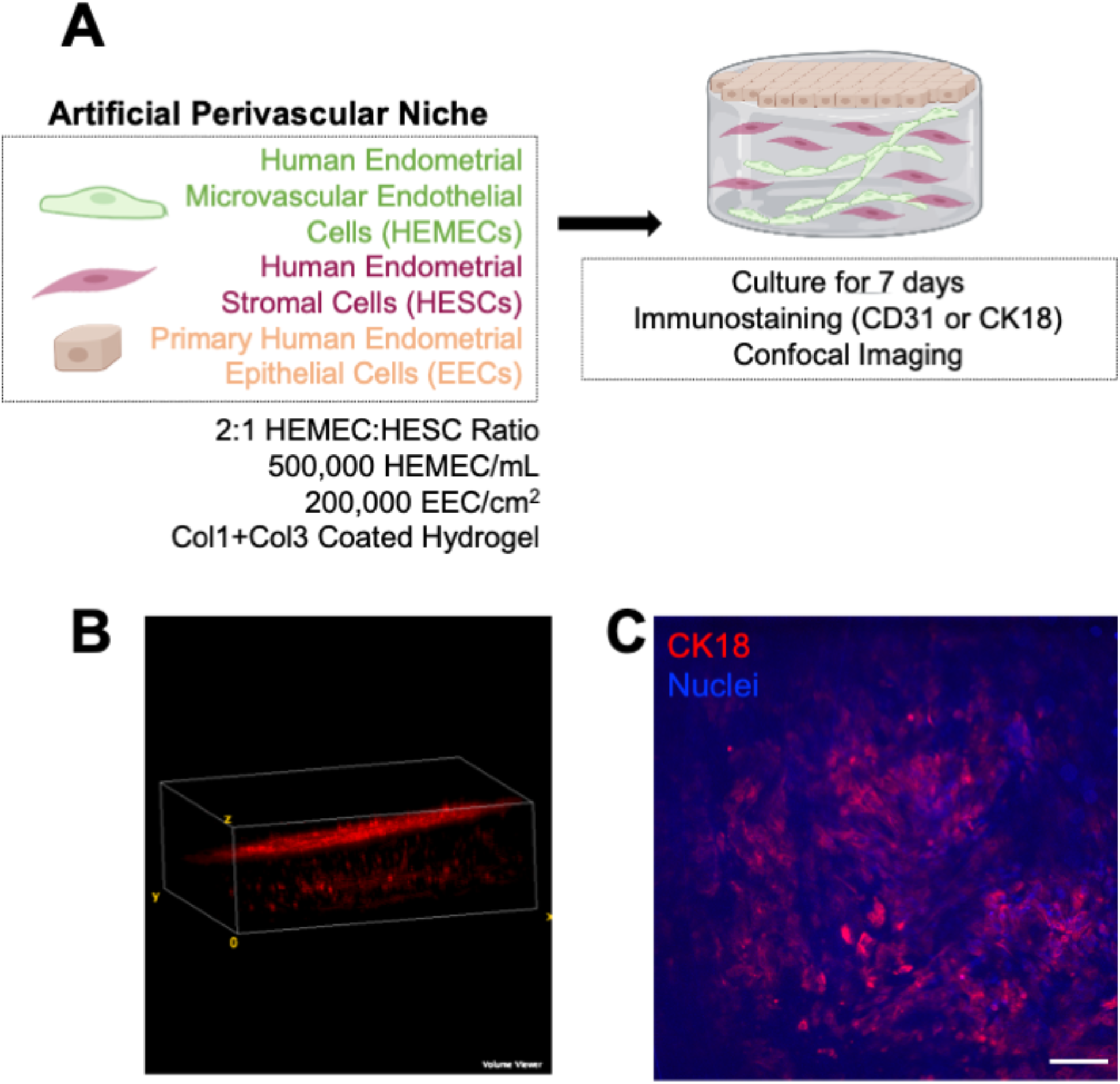
Fabrication of a stratified endometrial model. **(A)** Experimental summary. **(B)** FIJI Volume Viewer maximum intensity projection showing phalloidin-stained cells with EEC layer overlaying perivascular compartment. **(C)** Maximum intensity projections of Z-stacks of triculture hydrogel cultures stained for cytokeratin 18 (CK18). Scale bars: 100 μm. Created with Biorender.com.

## Discussion

Angiogenesis and vessel remodeling in the endometrium occurs during the menstrual cycle as the tissue is rebuilt and differentiates to prepare for potential pregnancy. Extensive remodeling of the existing vasculature occurs in response to infiltration of trophoblast cells from the blastocyst in order to provide blood flow to the growing fetus and placenta. Here, we report creation of a three-dimensional artificial endometrial perivascular niche embedded in a gelatin methacrylol hydrogel. 3D perivascular niche cultures demonstrate the capacity to exhibit morphological and biomolecular signatures of decidualization in response to exogenous hormones. These endometrial perivascular cultures provide a platform to examine reciprocal crosstalk between the perivascular niche and trophoblast cells. Further, the engineered endometrial perivascular niche hydrogel can be combined with our recently reported methods for endometrial epithelial cell culture to establish a stratified endometrial model consisting of primary endometrial epithelial cells overlaying the perivascular niche (*21*). Combined, these elements of a stratified endometrial model offers significant potential to gain mechanistic insight into endometrial remodeling and changes in endometrial vascular networks in response to decidualization.

Most existing *in vitro* models of vasculature utilize human umbilical vein endothelial cells (HUVECs) as their endothelial cell source due to their wide availability, capacity to be passaged numerous times, and excellent ability to form vessels *in vitro* (*22*). However, HUVECs are derived from umbilical cords and are not tissue-specific which calls into question the use of such cells for the development of tissue-specific vasculature models. Here, we utilize HEMECs as an endothelial cell source to create an endometrial-specific model of the endometrial perivascular niche. We also used human endometrial stromal cells (HESCs) as an endometrial stromal cell source. The addition of a stromal population to endothelial cells encourages and supports formation of endothelial networks for long-term culture (*23–25*). Without stromal cells, endothelial cells form endothelial structures that last only transiently and fall apart over time, consistent with our results from our tube formation assay. We identified 2:1 ratio of endothelial to stromal cells allowed for 20+ days of culture. Our studies suggest that reduced initial stromal cell density may be more conducive to perivascular stability, likely due to the contractile nature of stromal cells which can cause the hydrogel cultures to contract and disintegrate over time, a result consistent with other reports in the literature (*26, 27*).

We then demonstrate that endometrial perivascular cultures do not require exogenous pro-angiogenic factors (VEGF) to promote network formation. Endometrial cultures also express markers of vessel network maturity such as basement membrane protein deposition and tight junction markers. Our artificial perivascular cultures demonstrate similarities to the native endometrium. The average vessel length per branch point was determined to be approximately 100-200 μm across the menstrual cycle and this value varies depending on cycle phase (*28*). Although our average branch values were less than these *in vivo* values, we observed significant differences in network complexity with the addition of hormones. This indicates that perivascular cultures formed *in vitro* from endometrial derived endothelial and stromal cells are hormone responsive. Given emerging literature seeking to better define population variation in hormone concentration across the menstrual cycle, an engineered endometrial perivascular network may provide the opportunity to quantify shifts in vessel complexity in response to exogenous hormone signals representative of discrete menstrual cycle phases. Future work to assess lumen formation and patent vessels would allow for the creation of perfusable networks to assess molecule transport and vessel perfusion in the context of pregnancy.

We subsequently sought to determine how decidualization status of stromal cells affects the perivascular niche. Decidualization is necessary and critical to prepare the endometrium for a potential pregnancy (*3*). Recapitulating such processes may be important for an endometrial model system used to study implantation events. For these studies, we chose two decidualization protocols commonly used in the literature (*15–20*). One employs the synthetic progestin medroxyprogesterone acetate (MPA) and the other uses progesterone (P4). Across our studies, we observed differences in the effects of these two protocols in our perivascular cultures, notably the effects of the P4-decidualized condition did not seem to increase total network length/mm^3^, total vessels, and total branches as much as the MPA-decidualized condition. Analysis of our perivascular cultures in the absence and presence of decidualization hormones revealed an increasing trend for total network length/mm^3^, total number of branches, and total number of vessels with the addition of hormones. Interestingly, branch length significantly decreased with the addition of hormones. Observations from human specimens demonstrate that the endometrial spiral arterioles grow, lengthen, and coil during the secretory phase of the menstrual cycle (phase when decidualization occurs) (*1*). These data are consistent with our observations, further demonstrating that this model system captures endometrial physiologic responses *in vitro*.

Next, we analyzed shifts in the perivascular secretome in response to decidualization. Our analysis of the perivascular secreted factors detected 14 cytokines with statistically significant differences between the conditions: Activin A (Gene ID 3624), Amphiregulin (AR; Gene ID 374), Angiogenin (ANG; Gene ID 283), Angiopoeitin-1 (Ang-1; Gene ID 284), Endoglin (ENG; CD105; Gene ID 2022), Endostatin/Collagen XVIII (Gene ID 80781), Endothelin-1 (ET-1; Gene ID 1906), FGF acidic (FGF-1; Gene ID 2246), IGFBP-2 (Gene ID 3485), PDGF-AA (Gene ID 5154), Pentraxin 3 (PTX3; TSG-14; Gene ID 5806), Platelet Factor 4 (PF4; CXCL4; Gene ID 5196), Prolactin (Gene ID 5617), and Serpin F1 (PEDF; Gene ID 5176). These 14 proteins can be broadly characterized into proteins associated with angiogenesis, vessel stabilization, and vessel maturation (Angiogenin, Angiopoietin-1, Endostatin/Collagen XVIII, FGF-1, PDGF, PTX3, Serpin F1), proteins relevant to endometrial function and receptivity (Amphiregulin, Endoglin, Endothelin-1, IGFBP-2, PF4); and, proteins relevant to decidualization and stromal cells (Activin A, Angiogenin, PDGF, Prolactin).

Angiogenin, Angiopoietin-1, Endostatin/Collagen XVIII, FGF-1, PDGF, PTX3, and Serpin F1 are related to angiogenesis, vessel stabilization, and vessel maturation. The overall presence and corresponding changes in these factors in non-decidualized vs. decidualized conditions indicate that the perivascular cultures mature and stabilize over time and decidualization hormones directly impact endometrial angiogenesis. For example, low levels of Ang-1 are found in endometrial stromal fibroblasts and this decreases over the menstrual cycle (*29*). We observed a significant decrease in Ang-1 in one of the decidualization conditions compared to control. Although the second condition did not show a significant decrease, it did appear to be slightly lower than the control values. We also observed increases in endostatin and FGF-1 in decidualized conditions which could indicate vessel maturation, stabilization, and angiogenesis (*29, 30*)(*31*). Additionally, decidualization may induce proliferation or motility of cells because these conditions demonstrated increased expression of PDGF-AA, a growth factor shown to have roles in cell proliferation, angiogenesis, inflammation, and tissue repair (*32*).

Amphiregulin, Endoglin, Endothelin-1, IGFBP-2, PF4 are proteins relevant to endometrial function and receptivity. The secretion of these factors suggests our perivascular cultures demonstrate relevant endometrial cell behavior. For example, amphiregulin and CXCL4 were both shown to have increased secretion in decidualized samples. Amphiregulin is a member of the epidermal growth factor (EGF) family and has a role in uterine receptivity and blastocyst attachment (*33*). Amphiregulin has been found in the luminal epithelium at the site of blastocyst apposition and its expression is correlated with an increase in progesterone levels and blastocyst attachment (*33*). Our data also demonstrate this trend, with significantly increased levels of Amphiregulin in decidualized cultures containing increased progesterone. CXCL4 is also regulated by progesterone withdrawal which suggests it likely has a role in endometrial repair following menses (*34*). Our results were consistent with these data: CXCL4 secretion was increased in decidualized samples.

Activin A, Angiogenin, PDGF, Prolactin are proteins relevant to decidualization and stromal cells. Activin A is produced in high concentrations by decidualized stromal cells and interacts with matrix metalloproteinases (MMPs) to promote matrix remodeling in the decidual response (*35, 36*). Our data were consistent with these previous observations: we observed significantly increased levels of Activin A in decidualized perivascular hydrogels compared to control.

Although much of our data are consistent with the literature, differences could be due to donor variability or the use of cell lines instead of primary cells (*37–39*). For example, previous data using endometrial epithelial cells noted donor to donor differences in epithelial cell behavior so we would suspect to see potential differences in the use of donor-derived HEMECs as well (*37*). This could be further explored using additional cells from more donors. Furthermore, signaling from other cells in the endometrium (e.g., epithelial cells, natural killer cells, immune cells, etc.) could alter the secretome of stromal and endothelial cells which could account for some of these differences (*2, 40–42*). Expanded studies using additional endometrial cell types could begin to probe these differences and glean additional insights into the secretome of other endometrial cells.

We subsequently assessed crosstalk between the endometrial perivascular niche and Swan 71 trophoblast cells. The conditioned media from non-decidualized perivascular cultures increased trophoblast motility compared to the other tested conditions. Interestingly, conditioned media taken from the two decidualization cultures (MPA- and P4-decidualized) induced differential responses regarding trophoblast motility. Conditioned media from the P4-decidualized perivascular cultures increased trophoblast motility compared to the non-decidualized control (Swan71 invasion medium); however, the MPA-decidualized conditioned media induced less outgrowth compared to the non-decidualized control. Critically, these data show that endometrial perivascular niche decidualization status influences the activity of trophoblast cells via secreted factors. Although conditioned media (MPA- and P4-decidualized) from decidualized perivascular networks only induced a moderate increase in trophoblast motility, this only represents a single condition of hormone stimulation targeting initial decidualization events. There is a significant opportunity for future efforts to examine how trophoblast motility changes in response to a dynamic secretome based on hormone concentrations representative of greater shifts across the menstrual cycle. Notably, treatment of perivascular cultures with Swan71 trophoblast conditioned medium did not change the majority of markers used to assess vessel complexity; however, we did observe decreased branch length in the media control condition (50:50 ratio Swan71 invasion medium and endothelial growth medium) compared to the control condition (Swan71 invasion medium). This could be because the Swan71 unconditioned growth medium contains less proangiogenic factors compared to endothelial growth media. Interestingly, the signaling was not bi-directional. Although endometrial perivascular niche decidualization status influences the activity of trophoblast cells, trophoblast cells do not appear to influence structural changes in the endometrial perivascular niche cultures. This finding has interesting implications as to the control of early stages of trophoblast motility from the maternal endometrium but no marked early changes in the maternal perivascular architecture in response to initial trophoblast implantation. These findings suggest that the endometrium may have biological agency in the uterine-trophoblast interfacing process during implantation; this is a fascinating topic that is worthy of more in depth studies.

Finally, we demonstrate the creation of an endometrial triculture consisting of an endometrial epithelial layer overlaying the embedded perivascular system. Our work herein demonstrates an endometrial model of increased complexity compared to existing models that incorporates three endometrial cell types in one model system. We chose collagen I and collagen III as the basement membrane layer as our prior work demonstrated this combination of ECM biomolecules resulted in the best epithelial cell attachment (*43*). Our model expands upon existing stratified endometrial model systems because we have added additional complexity by not only including stromal cells but also endothelial cells to create an embedded vascular niche rather than only an embedded stroma (*37*).

Future work will focus on quantification of vessel network metrics for the triculture as well as on the hormonal work to determine how decidualization affects not only the perivascular compartment but also the epithelial layer. Additional opportunities for these studies include development an endometrial perivascular niche using primary endometrial stromal cells. HESCs are the most widely used cell line for endometrial stromal cells; however, as hTERT-immortalized cells, they may not mimic endometrial stromal cells as closely as primary cells could. The use of patient-derived cells could ameliorate this challenge and provide additional insights into endometrial perivascular function. Additionally, this work considers sex steroid hormone profiles from one point in the menstrual cycle. Ongoing work is looking to quantify metrics of network formation across the entire menstrual cycle to assess cyclic vessel formation and remodeling. As studies in humans have shown variation in menstrual cycle length and hormone profiles (*44, 45*), there is significant opportunity to use this platform as a route to explore patient variation via incorporation of different hormone profiles.

In conclusion, we describe the creation of an artificial endometrial perivascular niche embedded in GelMA hydrogels. Engineered endometrial perivascular cultures display hormone-responsiveness in our cultures, including variation in network complexity and secretion of soluble factors. We show a model of unidirectional signaling; although perivascular network conditioned medium increased trophoblast motility in spheroid motility assays, trophoblast conditioned medium showed limited effect on perivascular niche complexity. Finally, we describe a stratified endometrial model consisting of an endometrial epithelium overlaying an embedded perivascular niche. Tissue engineering models such as these not only provide novel platforms for assessing endometrial function but also allow us to probe questions regarding implantation that are currently impossible to answer in humans due to ethical constraints, challenging time points, and lack of imaging modalities. With the creation of these platforms, we hope to provide researchers with novel technologies that can further the field of uterine health.

## Materials and Methods

### Experimental Design

The objective of this study was to design, characterize, and implement and artificial tissue engineered model of endometrial vasculature. Using gelatin methacrylol hydrogels, we create a co-culture system of endometrial endothelial and stromal cells and subsequently assess response to sex steroid hormones and trophoblast secreted factors.

### Cell Culture and Maintenance

#### Human Endometrial Microvascular Endothelial Cell Culture

Human endometrial microvascular endothelial cells (HEMEC; ScienCell #7010) were maintained as per the manufacturer’s instructions in phenol red-free Endothelial Cell Medium (ECM; ScienCell #1001-prf) supplemented with an endothelial cell growth supplement (ScienCell #1052), 5% charcoal-stripped fetal bovine serum (Sigma-Aldrich F6765), and 1% penicillin/streptomycin (ThermoFisher 15140122). Charcoal-stripped fetal bovine serum was used to reduce the steroid hormone concentrations in the cell medium. HEMECs were cultured on bovine plasma fibronectin (ScienCell #8248) coated vessels. HEMECs were used experimentally no more than 5 passages from purchase. HEMECs were cultured in 5% CO_2_ incubators at 37°C. Routine mycoplasma testing was performed using a MycoAlert™ Mycoplasma Detection Kit (Lonza). Cell ancestry information (e.g., racial and ethnic background, age, gender identity) was not provided by the vendor although the cell ancestry may affect cellular behavior and response (*38, 39*).

#### Human Endometrial Stromal Cell Culture

Human endometrial stromal cells (HESC; ATCC® CRL-4003) were maintained as per the manufacturer’s instructions in custom phenol red-free DMEM/F-12 (based on Sigma #D 2906) supplemented with 1% ITS+ Premix (Corning 354352), 500 ng/mL puromycin (Millipore Sigma P8833), 10% charcoal stripped fetal bovine serum (Sigma-Aldrich F6765), and 1% penicillin/streptomycin. HESC were used experimentally no more than 5 passages from purchase. HESC were cultured in 5% CO_2_ incubators at 37°C. Routine mycoplasma testing was performed using a MycoAlert™ Mycoplasma Detection Kit (Lonza). Cell ancestry information (e.g., racial and ethnic background, age, gender identity) was not provided by the vendor although the cell ancestry may affect cellular behavior and response (*38, 39*).

#### Primary Human Endometrial Epithelial Cell Culture

We cultured primary human endometrial epithelial cells (EECs; LifeLine Cell Technology FC-0078; Lot 03839; Caucasian Female Donor, 33 y.o., uterine prolapse) as per the manufacturer’s instructions in phenol red-free medium (LifeLine Cell Technology) and in 5% CO2 incubators at 37°C. EECs were used experimentally at two passages from receipt. Cells were routinely tested for mycoplasma contamination using the MycoAlertTM Mycoplasma Detection Kit (Lonza).

#### 2D Culture of Human Endometrial Microvascular Endothelial Cells

HEMEC cells were seeded into individual wells of a 6 well plate and cultured until confluent. Cells were then fixed in formalin (Sigma-Aldrich), permeabilized for 15 minutes in 0.5% Tween20 (Fisher Scientific BP337), washed 3×5 minutes with 0.1% Tween20 solution (PBST), blocked with 2% Abdil (2% bovine serum albumin; Sigma Aldrich A4503 + 0.1% Tween20) for 1 hour, and stained with primary antibodies (1:200 CD31; Dako IS610 or 2.5 μg/mL von Willebrand Factor viii; Invitrogen MA5-14029) overnight at 4°C. 5×5 minute PBST washes were performed followed by staining with secondary antibody (1:500 Alexafluor 488 goat anti-mouse; Thermo Fisher A-11001) overnight at 4°C. 5×5 minute PBST washes were performed followed by staining with Hoechst (1:2000; Thermo Fisher H3570) for 10 minutes at room temperature. One final PBST wash was performed and cells were stored in PBST until imaged. Wells were imaged using a Leica DMI 4000 B Microscope (Leica Microsystems).

#### Matrigel Tube Formation Assay

100 μL of phenol red-free Matrigel (1.35 mg protein/well; Corning 356237) was pipetted into each well of a 96 well plate and polymerized in the incubator. 10,000 HEMECs were added per well (n=8 wells). Each well was imaged at 6 hours and 12 hours after seeding using a Leica DMI 4000 B Microscope (Leica Microsystems).

### Synthesis and Fabrication of Methacrylamide-Functionalized Gelatin (GelMA) Hydrogels

GelMA was synthesized, dialyzed, lyophilized, and was found to have a degree of functionalization of 57%, determined via ^1^H-NMR (*46–48*). Prior to cell culture experiments, lyophilized GelMA was sterilized for 30 minutes under UV light. Hydrogels were fabricated using a solution consisting of lyophilized GelMA (5 wt%) dissolved at 37°C in phosphate buffered saline (PBS; Lonza 17-516F) and combined with 0.1% w/v lithium acylphosphinate (LAP) as a photoinitiator. Hydrogels were polymerized under UV light (λ=365 nm, 7.14 mW cm^-2^; AccuCure Spot System ULM-3-365) for 30 s.

### Endometrial Perivascular Niche Hydrogel Co-Cultures

#### Co-culture Fabrication and Maintenance

HEMEC and HESC were passaged and encapsulated in GelMA hydrogels at 1:1, 1:2, and 2:1 endothelial:stromal cell ratios. The concentration of endothelial cells was kept consistent with each ratio at 500,000 HEMEC/mL and the concentration of stromal cells was calculated from this value and the ratios. Hydrogels were cultured in 48 well plates for 7 days and maintained in ECM with or without additional growth factors (± 100 ng/mL recombinant human VEGF_165_; PeproTech 100-20) and hormones. The medium for hydrogel samples was replaced every 3 days (800 μL/well). The endogenous VEGF concentration in ECM was reported to be 2 ng/mL by the vendor (ScienCell). All experiments except those in Fig. 2, 3, and 4 used charcoal-stripped fetal bovine serum (Sigma-Aldrich F6765) instead of regular fetal bovine serum to decrease endogenous hormones in the base medium.

#### Co-culture Decidualization

Decidualization of endometrial stromal cells was induced by culturing hydrogels in the presence of the following decidualization hormone cocktails: (i) based on synthetic progesterone, 1 μM medroxyprogesterone acetate (MPA; Sigma-Aldrich M1629) + 0.5 mM 8-bromodenosine 3’,5’-cyclic monophosphate (8-Br-cAMP; Sigma-Aldrich B5386) or (ii) progesterone based, 0.5 mM dibutyryl cyclic AMP (dcAMP; Millipore Sigma 28745) + 10 nM estradiol (E2; Sigma-Aldrich E2758) + 100 nM progesterone (P4; Sigma-Aldrich P8783). Control samples had no hormones added to the medium. Medium was replaced every 3 days, collected at days 3 and 6, and stored at −80°C.

### Characterization of Perivascular Niche Cultures

#### Immunofluorescent Staining

On day 7 of culture, hydrogel samples were fixed with formalin (Sigma-Aldrich) and washed three times with PBS. Hydrogels were permeabilized for 15 minutes in a 0.5% Tween20 (Fisher Scientific BP337) solution and washed 3×5 minutes in 0.1% Tween20 solution (PBST). Samples were blocked for 1 hour at room temperature in a 2% Abdil solution (2% bovine serum albumin; Sigma Aldrich A4503 + 0.1% Tween20) and subsequently incubated in primary antibody solution (1:200 CD31 Dako IS610 + 1:200 CD10 Invitrogen PA5-85875 or 1:200 anti-laminin Abcam ab11575 or 5 μg/mL ZO-1 Invitrogen #61-7300) overnight at 4°C. 4×20 minutes washes with PBST were performed and then cultured in secondary antibody (1:500 Alexafluor 555 goat anti-rabbit Thermo Fisher A-21428 and/or 1:500 Alexafluor 488 goat anti-mouse Thermo Fisher A-11001) overnight at 4°C. Hydrogels were washed 4×20 minutes with PBST and then incubated for 30 minutes in Hoechst (1:2000; Thermo Fisher H3570). Samples were washed a final time in PBST and were stored in PBST until imaged.

#### Microscopy Techniques

Hydrogels were imaged using glass bottom confocal (In Vitro Scientific, D29-20-1-N) dishes on a DMi8 Yokogawa W1 spinning disc confocal microscope outfitted with a Hamamatsu EM-CCD digital camera (Leica Microsystems). Three 100 μm z-stacks with a 5 μm step size were taken for each hydrogel for 3 regions of interest (ROI) except for time course experiments and laminin/ZO-1 stained hydrogels (n=2-3 hydrogels; n=2 ROI per hydrogel). For day 14 and day 21 hydrogels, 1 ROI was imaged which captured roughly 80-100% of the entire gel area. Fluorescent images were artificially brightened for figures but not for analysis.

#### Image Analysis

Images were analyzed using a computational pipeline consisting of a FIJI macro and custom MATLAB algorithm (*49, 50*). This pipeline allows for 3D quantification of vessel networks across image z-stacks. Briefly, individual images of each z-stack were blurred, filtered, and binarized using a FIJI macro. Then, the binarized images were analyzed using a custom MATLAB algorithm that quantified total branches + endpoints, branch points, number of vessels, and total network length from skeletonized images. Using Microsoft Excel, we then calculated the total network length / mm^3^, average branch length (network length / number of vessels), number of branches, and number of vessels for each sample.

To compute the degree of overlap between CD31 signal and laminin/ZO-1 signal, CD31 and laminin/ZO-1 Z-stacks were binarized using the same FIJI macro listed above. Compressed copies of the Z-stacks were also created to match the size of the binarized images. Average pixel intensity was generated for both stains and the degree of overlap was then calculated by multiplying the binarized matrix of vessels by the binarized matrix of the proteins (Unpublished method from Victoria Barnhouse et al. *in preparation*).

#### Cytokine Array

A Proteome Profiler Human Angiogenesis Array (R&D Systems ARY007) was used to determine relative levels of 55 angiogenesis-related proteins. Medium was collected at days 3 and 6 of culture, stored at −80°C until use, and pooled for analysis. 500 μL of medium was used for each day (1 mL total per cytokine array; n=3 samples per condition).

The array was run as per the manufacturer’s instructions and imaged (4 minute exposure) using an Amersham ImageQuant 800 Fluor system (Cytiva). Pixel density of each array spot was quantified using FIJI. The negative control spot averages were subtracted from the pixel density of each sample and then pixel density of each sample was normalized to the pixel density of positive control spots.

#### STRING Analysis

Statistical analysis was performed to determine which of the 55 angiogenesis-related proteins were statistically significantly different between groups. The resultant analysis revealed 14 significant proteins. These 14 proteins were entered into the STRING (Search Tool for the Retrieval of Interacting Genes/Proteins) Database to determine known and predicted protein-protein interactions (*51–53*). A network summary view was created using a medium confidence minimum required interaction score (0.400).

### Trophoblast and Perivascular Niche Interactions

#### Perivascular Hydrogel Conditioned Media Effects on Trophoblast Motility

Control and decidualized perivascular hydrogels were cultured as described above. Media were collected during media changes on days 3 and 6 of culture, filtered, and stored at −20°C until use. Unconditioned ECM was also collected for use as a control. Conditioned media from days 3 and 6 were pooled prior to adding to spheroid cultures. Spheroid motility assays were performed as described previously by our group (*47, 48, 54*). For these studies, we used Swan 71 cells derived from a 7-week first trimester placenta; however, no additional donor information was provided (*55*). Swan71 at passage one from receipt were cultured in growth medium consisting of phenol red-free DMEM (SCS Cell Media Facility, UIUC) supplemented with 10% charcoal-stripped fetal bovine serum, 1% penicillin/streptomycin, and 500 ng/mL puromycin. Once passaged for experiments, the cells were cultured in phenol red-free DMEM, 2% charcoal-stripped fetal bovine serum, and 1% penicillin/streptomycin (Swan71 motility medium). Cells were cultured in flasks until 80-90% confluence and added to round bottom plates (Corning 4515; 4,000 cells/well) for at least 48 hours on a shaker (60 rpm) in the incubator until spheroids formed. Individual spheroids were encapsulated in GelMA hydrogels and maintained in 800 μL of medium (Swan71 motility medium, 50:50 Swan71 motility medium:ECM, or 50:50 Swan71 motility medium:PVN-conditioned medium) for 3 days. Each encapsulated spheroid was imaged daily on a Leica DMI 4000 B microscope (Leica Microsystems). Total outgrowth area was calculated using the measure tool in FIJI by averaging three traced measurements of the outgrowth area. Fold change was calculated by normalizing outgrowth area to initial spheroid area (day 0).

#### Trophoblast Conditioned Media Effects on Perivascular Hydrogels

Swan71 at passage one from receipt were cultured in growth medium consisting of phenol red-free DMEM (SCS Cell Media Facility, UIUC) supplemented with 10% charcoal-stripped fetal bovine serum, 1% penicillin/streptomycin, and 500 ng/mL puromycin. Swan71 cells were cultured in a T75 culture flask and medium was collected at confluence, syringe filtered, and stored at −20°C until use. Unconditioned growth medium was also collected for use as a control. Co-culture hydrogels were fabricated and cultured as described above in ECM (control), a 50:50 ratio of ECM to unconditioned Swan71 growth medium (media control), or 50:50 ratio ECM to Swan71 conditioned medium. Hydrogels were stained, imaged, and analyzed as described above.

### Triculture of Endometrial Endothelial, Stromal, and Epithelial Cells

Perivascular hydrogel cultures were prepared as described above and cast into Ibidi μ-Slides Angiogenesis (10 μL prepolymer solution; Ibidi 81506). Polymerized gels were then coated with Collagen 1 (EMD Millipore 08-115MI) and Collagen 3 (EMD Millipore CC054) using microbial transglutaminase (mTg; Zedira T001) (*43, 56, 57*). A 1:1 ratio of 0.5 mg/mL mTg and 10 μg/mL ECM protein (1:1 ratio Collagen 1 and Collagen 3) were combined and 20 μL of this solution was pipetted onto hydrogels. Coated hydrogels were incubated for 1 hour in 5% CO2 incubators at 37°C. A quick wash was performed using 20 μL of PBS. After the wash step, we seeded 200,000 EEC/cm^2^ onto hydrogels. We cultured tricultures for 7 days and subsequently stained them with CD31 and phalloidin (7 μL per 1000 μL solution) or cytokeratin 18 (CK18; 1:250, Cell Signaling 24E10) using the protocol described above. We then took Z-stack images of each gel from the top of the gel as far down as we could visualize. We took 1 Z-stack per gel (n=2 gels per condition).

### Statistical Analysis

OriginLab 2021b and RStudio were used for statistical analyses. Normality was determined via Shapiro-Wilkes and homoscedasticity was determined via Levene’s test. Data were analyzed using a one-way analysis of variance (ANOVA) and Tukey post hoc test (normal, homoscedastic), Welch’s ANOVA and Games-Howell post hoc test (normal, heteroscedastic), Kruskall-Wallis ANOVA and Dunn’s post hoc test (non-normal, homoscedastic), or Welch’s Heteroscedastic F Test with Trimmed Means and Winsorized Variances and Games-Howell post hoc test (non-normal, heteroscedastic). Significance was set as *p*<0.05 and data are presented as mean ± standard deviation unless otherwise described. Each quantitative experiment used n=3-6 hydrogels unless otherwise noted. Plots were generated using OriginLab.

## Acknowledgments

The content herein is solely the responsibility of the authors and does not necessarily represent the official views of the National Institutes of Health. The authors also gratefully acknowledge additional funding provided by the Department of Chemical & Biomolecular Engineering and the Carl R. Woese Institute for Genomic Biology at the University of Illinois at Urbana-Champaign. The authors thank Drs. Gil More (Yale University School of Medicine, New Haven, CT) and Gabriela Dveksler (Uniformed Services University of Health Sciences, Bethesda, MD) for providing the Swan71 cells. The authors also thank the Institute for Genomic Biology Core Facilities (Dr. Austin Cyphersmith) at the University of Illinois Urbana-Champaign for assistance with imaging. The authors would like to also thank Dr. Sara Pedron Haba, Dr. Victoria Barnhouse, and Vasiliki Kolliopoulos for excellent technical assistance.

## Funding

Research reported was supported by:

National Institutes of Diabetes and Digestive and Kidney Diseases of the National Institutes of Health R01 DK0099528 (BACH)

National Cancer Institute of the National Institutes of Health R01 CA256481 (BACH)

National Institute of Biomedical Imaging and Bioengineering of the National Institutes of Health T32 EB019944 (SGZ)

Department of Chemical & Biomolecular Engineering University of Illinois Urbana-Champaign

Carl R. Woese Institute for Genomic Biology University of Illinois Urbana-Champaign.

## Author contributions

We describe contributions to the manuscript using the Contributor Roles Taxonomy (CRediT) (*58, 59*):

*Writing – Original Draft:* SGZ
*Writing – Review & Editing:* SGZ, HT, IJ, CC, JZ, NC, IP, KC, BACH
*Conceptualization:* SGZ and BACH
*Investigation:* SGZ
*Methodology:* SGZ
*Formal Analysis:* SGZ
*Data Curation:* SGZ
*Visualization:* SGZ
*Project Administration:* BACH
*Resources:* BACH;
*Funding Acquisition:* BACH
*Supervision*: BACH
*Miscellaneous:* HT and IP assisted with experimentation. IJ assisted with R Code. NC and IP assisted with literature review. CC and JZ provided the image analysis pipeline.

## Data and materials availability

The raw data required to reproduce these findings are available per request by contacting the corresponding author. The processed data required to reproduce these findings are available per request by contacting the corresponding author.

## Supplementary Materials

Supplemental information can be found in the Supplementary Materials.

### Supplementary Text

**Fig. S1.**
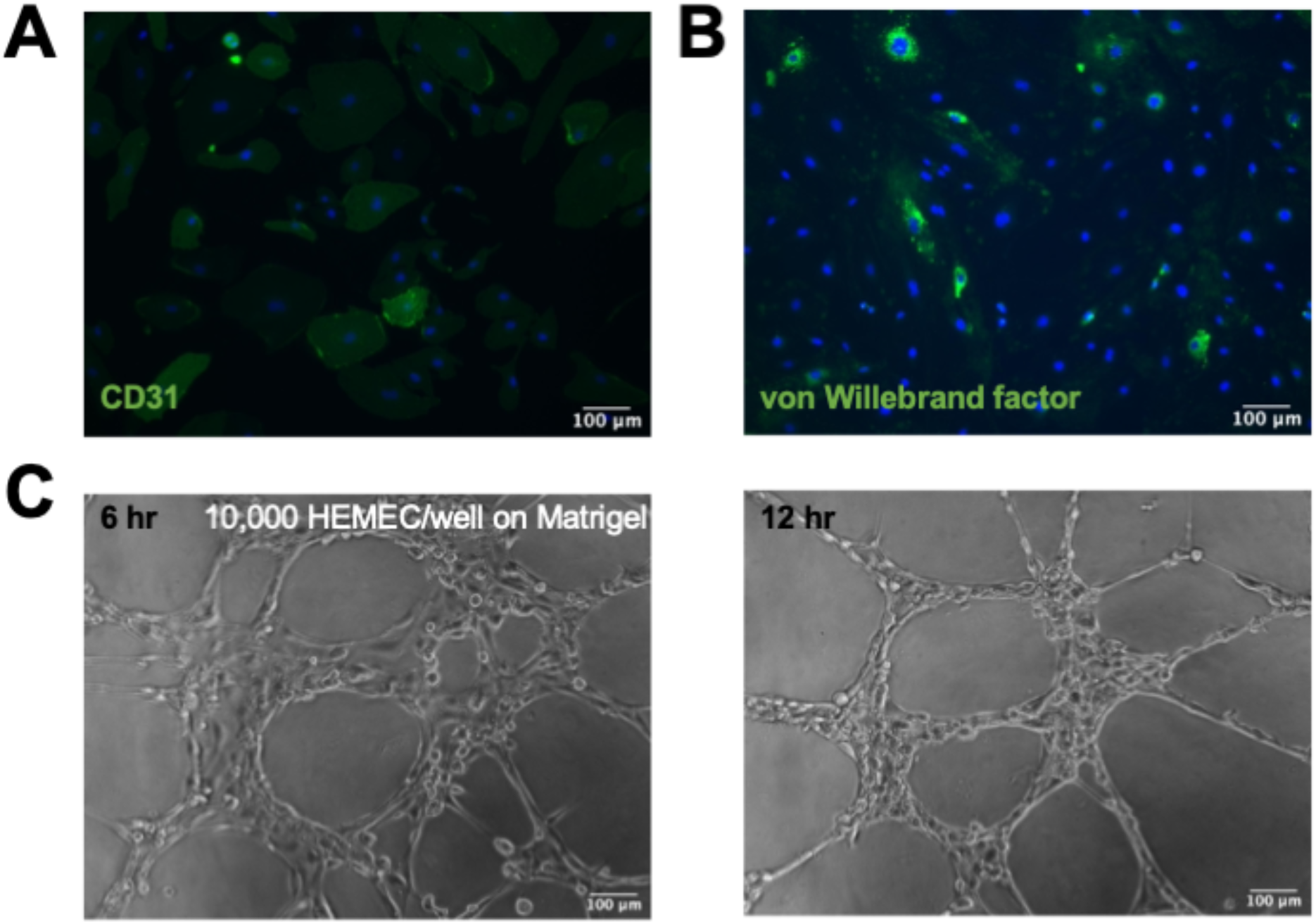
Angiogenic potential of human endometrial microvascular endothelial cells (HEMEC). HEMEC cultured on well plates express characteristic endothelial cell markers such as (A) CD31 (cluster of differentiation 31) and (B) von Willebrand factor. (C) HEMEC demonstrate the ability to form tubes on Matrigel that form transiently and fall apart in less than 24 hours. Scale bar: 100 μm.

**Fig. S2.**
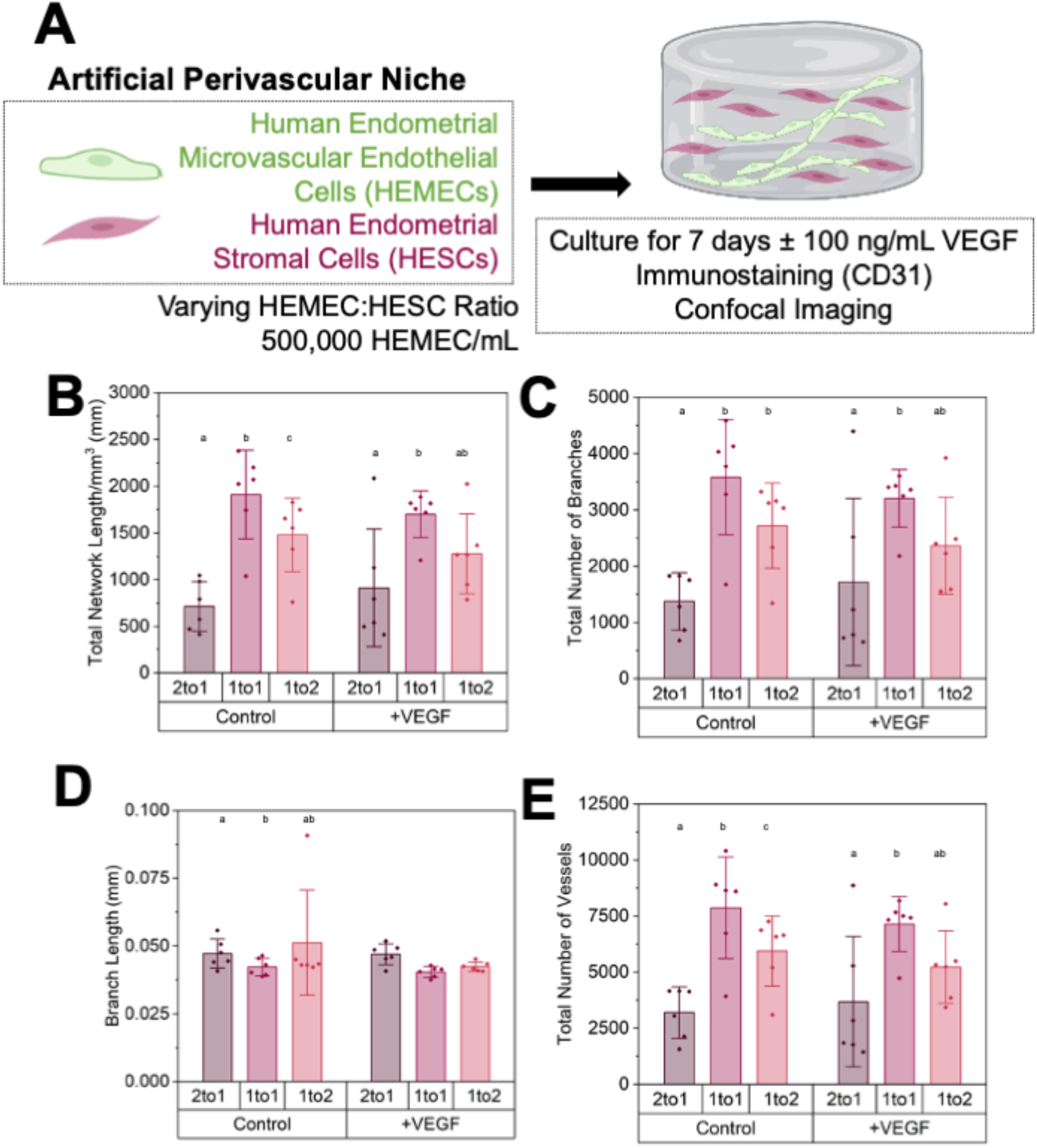
Optimization of an artificial endometrial perivascular niche. (A) Experimental summary. (B) Quantification of total vessel length per mm^3^, (C) total number of branches, (D) average branch length, and (E) total number of vessels for control and vascular endothelial growth factor (VEGF) samples (n=6 hydrogels per condition; 3 ROI imaged per gel and averaged) of varying endothelial to stromal cell ratios. Groups with different letters are statistically significantly different from each other. Data presented as mean ± standard deviation. Created with Biorender.com.

**Fig. S3.**
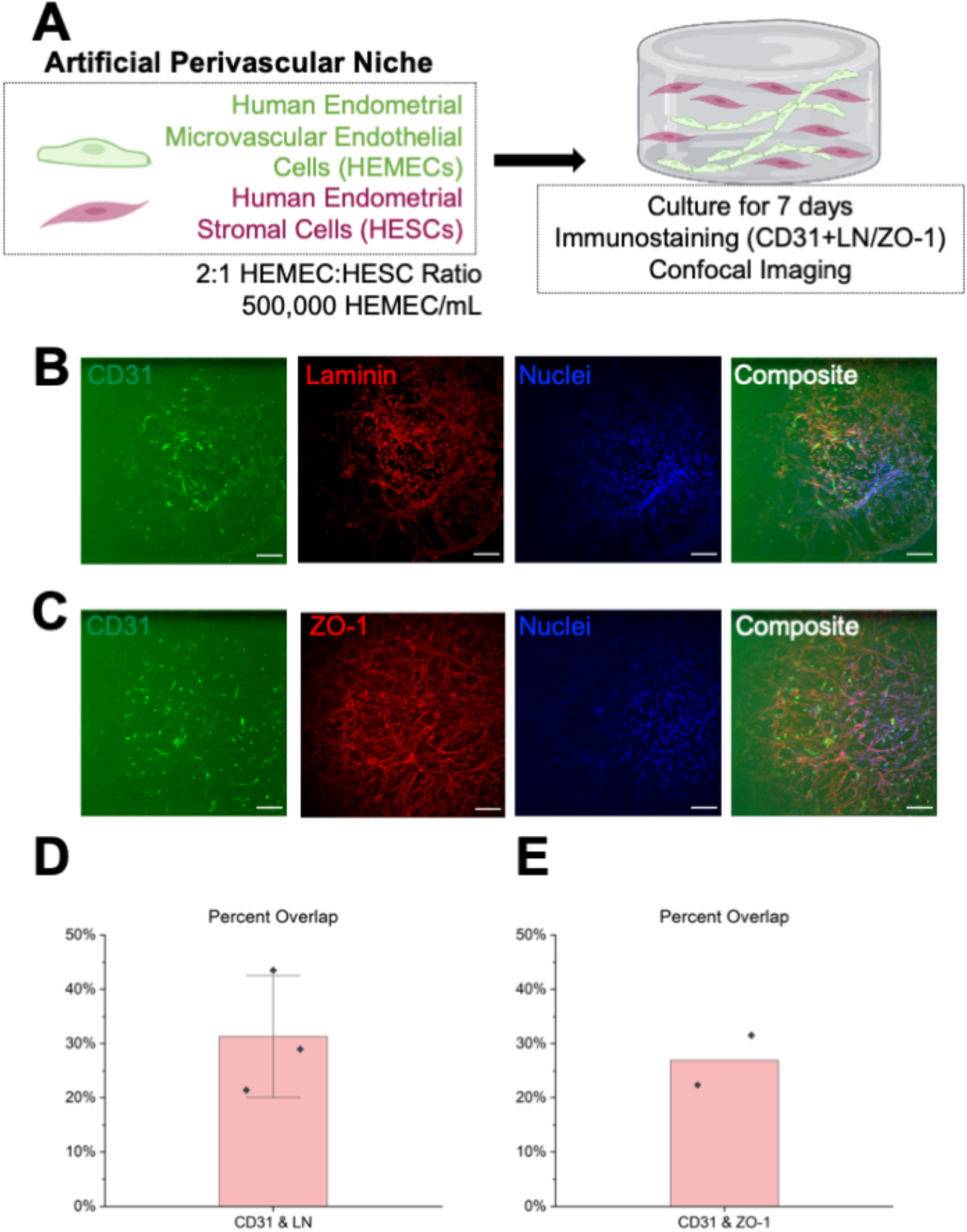
Characterization of extracellular matrix deposition and tight junction expression in perivascular cultures. (A) Experimental summary. Maximum intensity projections of Z-stacks of artificial endometrial perivascular niche hydrogel cultures stained for CD31 (HEMEC-endothelial cells) and (B) laminin (C) and ZO-1. Green-CD31; Red-Laminin or ZO-1; Blue-Nuclei. Percent overlap was calculated between CD31 signal and (D) laminin and (E) ZO-1. n=2-3 hydrogels per condition. Data presented as mean ± standard deviation. 2 ROI imaged per gel. Scale bars: 100 μm. Images artificially brightened for visualization using FIJI. Created with Biorender.com.

**Fig. S4.**
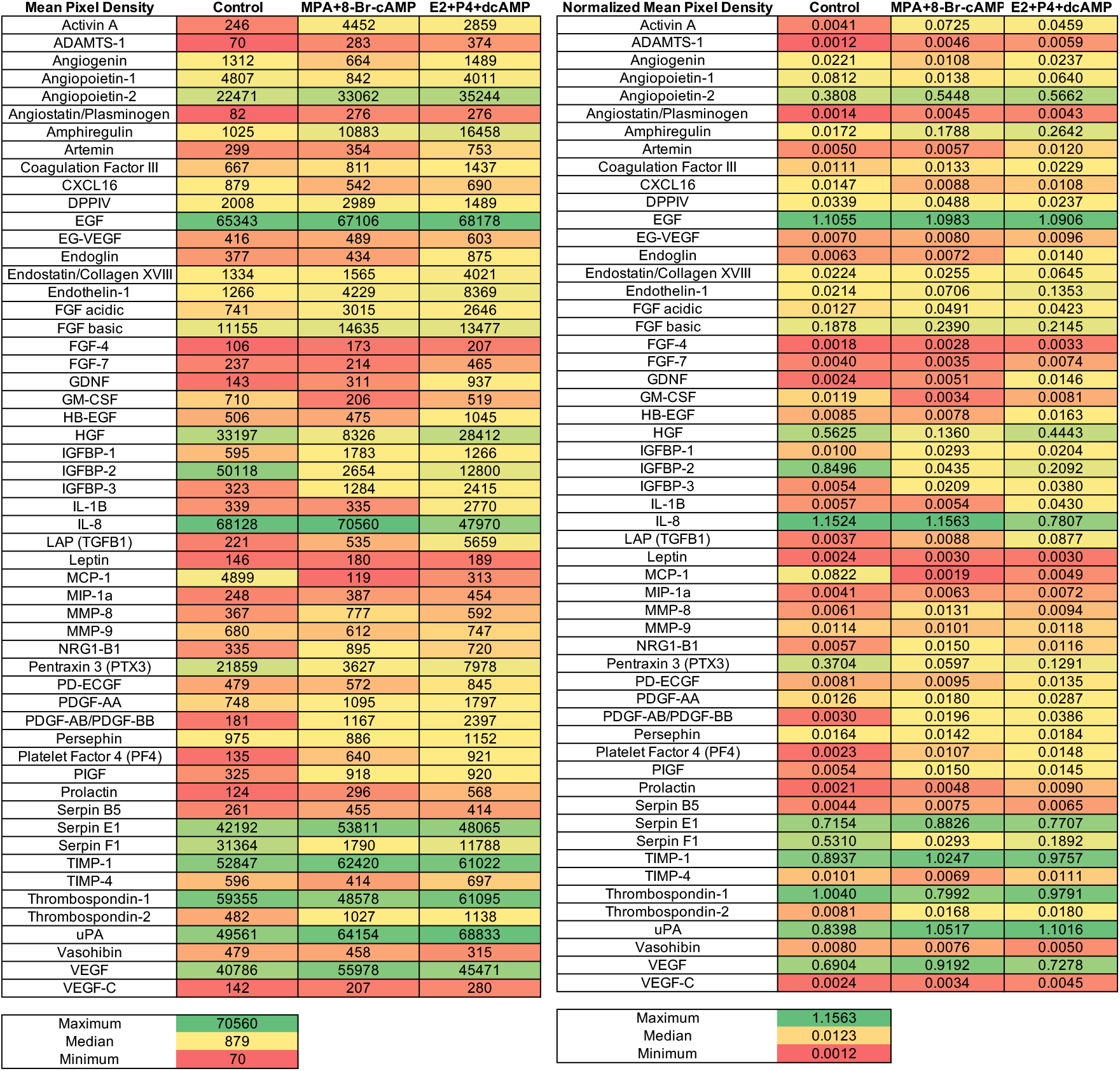
Cytokine array data. **A.** Raw mean pixel density values. **B.** Mean pixel density values normalized to positive control spots.

**Fig. S5.**
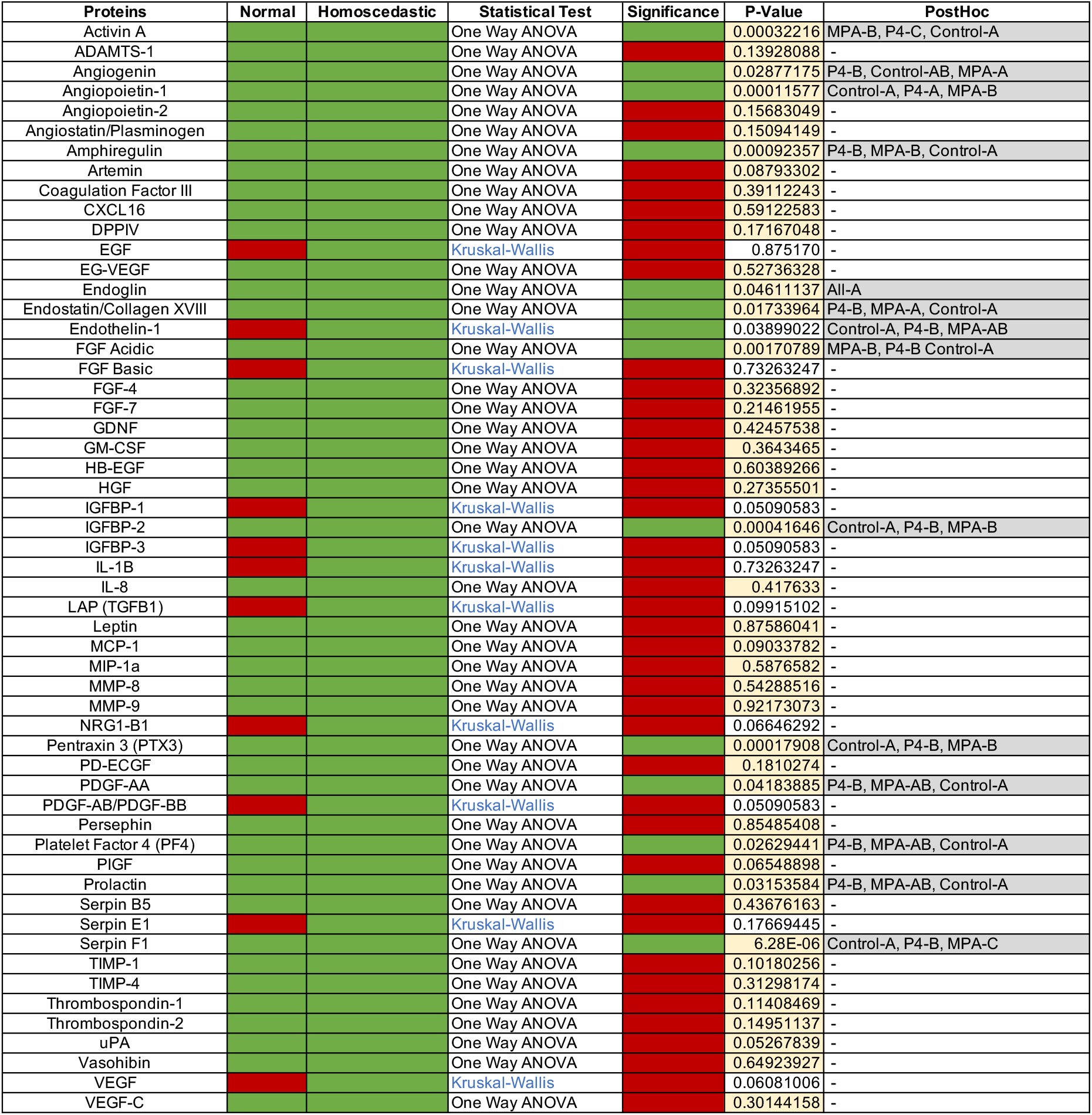
Statistical analysis of 55 cytokines. Green: Yes. Red: No.

